# Network Structure and Social Complexity in Primates

**DOI:** 10.1101/354068

**Authors:** R.I.M. Dunbar

**Affiliations:** Department of Experimental Psychology, University of Oxford, South Parks Road, Oxford OX1 3UD, UK; Department of Computer Science, Aalto University, FI-00076 Espoo, Finland [ ]

**Keywords:** social networks, grooming, multilevel sociality, time budgets, cognition

## Abstract

Primates use social grooming to create and maintain coalitions. Because of this, individuals focus their time on a small number of individuals, and this means that in many cases group networks are not fully connected. I use data on primate grooming networks to show that three different social grades can be differentiated in terms of network structuring. These grades seem to arise from a glass ceiling imposed on group size by limits on the time available for social grooming. It seems that certain genera have managed to circumvent this constraint by a phase shift in the behavioural and cognitive mechanisms that underpin social relationships in a way that allows a form of multilevel sociality based on weak and strong ties not unlike those found in human social networks.

## Introduction

In primates, social grooming is the principal process by which relationships are built up and serviced (Dunbar & Shultz 2010; Massen et al. 2010; Silk et al. 2010a,b). Although time devoted to grooming increases linearly with group size (Dunbar 1991; Lehmann et al. 2007; Dunbar & Lehmann 2013), this does not mean that individuals groom with every other member of the group; rather, it is a consequence of the fact that individuals choose to invest disproportionately in a small number of core relationships (Dunbar 1980; Duboscq et al. 2016), probably so as to ensure their functionality as coalitions (Dunbar 2012). In both primates (Dunbar 1980) and humans (Roberts & Dunbar 2011), the quality of a relationship is determined by the amount of time an individual invests in it (in the primate case, by grooming with its partner): grooming partners are more likely to come to each other’s aid in altercations (Dunbar 1980) and to alert when a partner gives fear screams (Seyfarth & Cheney 1984). More importantly, these relationships demonstrably influence female stress levels, fecundity and fitness in both primates (Silk et al. 2003, 2010a,b; Wittig et al. 2008) and equids (Cameron et al. 2009). If grooming relationships are the basis of social bonding in primates (Dunbar & Shultz 2010; Dunbar et al. 2009; Silk et al. 2010b), this raises questions about how group cohesion and stability are engineered in social groups that greatly exceed an individual’s capacity to groom others (Dunbar 2012). This seems to have been a question that, so far, no one has thought of asking.

I use comparative data on grooming networks in a taxonomically wide set of primates to determine how social structure varies with group size. I focus explicitly on grooming relationships, rather than affiliative behaviours like proximity that some recent studies (e.g. Pasquaretta et al. 2014) have used because grooming is the central engine in building relationships for primates: grooming triggers the release of endorphins in the brain in both primates (Fabre-Nys et al. 1982; Keverne et al. 1989; Martel et al. 1993) and humans (Nummenmaa et al. 2016), and endorphins are central to the creation and management of affiliative relationships (Panksepp 1999; Depue & Morrone-Strupinsky 2005; Dunbar 2010; Machin & Dunbar 2011; Pearce et al. 2017). Proximity and huddling merely imply tolerance (a consequence), whereas grooming, and especially its reciprocation, implies something about active engagement and commitment to another individual (a mechanism).

I focus on two indices of group social structure: the grooming clique (the number of other adults that an individual grooms with on a regular basis – in other words, its core coalition partners) and the grooming network (the number of individuals linked in a continuous chain of such relationships). The first tells us something about the social skills of an individual monkey (how many relationships it can keep functional at the same time). The second tells us something about the structural stability of a social group: groups that are split into several weakly connected networks may be more likely to fission when the ecological and social costs of group-living become intolerable (Dunbar et al. 2009).

I first examine the relationship between these two indices to determine how grooming clique size relates to network size and how network size relates to group size in order to explore the structural properties of groups. In both cases, we are interested in whether two indices form a single monotonic functional relationship or a set of separate but parallel grades. A single relationship in both cases tells us that these indices form a simple fractal pattern in which one layer scaffolds the next; living in large groups is thus a consequence of being able to manage more grooming relationships in the base layer, and nothing more. If the relationship between two indices also involves grades, such that there are two or more monotonic relationships involved, this would imply that some species are able to maintain a higher order grouping (an extended network) without necessarily having to change their lower level grouping (grooming clique size) proportionately; this implies an ability to manage two different kinds of relationships (weak and strong) simultaneously without necessarily having to invest equally in all of them. If so, we will want to know whether these grades coincide with any obvious correlated change in grooming behaviour or cognition. For present purposes, I will use neocortex volume and a direct measure of executive function competence as relevant indices for this.

## Methods

Data on the frequency of social grooming among adults are taken from Kudo & Dunbar (2001), still the most comprehensive grooming network dataset available, supplemented by three more recent studies. Pair-living species are excluded, since grooming networks are meaningless when there are only two adults. In total, 101 social groups drawn from 34 species are included in the sample, a larger and broader coverage than any previous analysis has used. The grooming data for each group are cast in a triangular matrix and used to calculate the two structural indices (grooming clique size for each individual and the mean size of all grooming networks in the group). In the network analysis literature, these indices are conventionally referred to as the *degree* and *n-clique*, respectively, but I will use the terms grooming clique and network here since they are more meaningful.

An important issue in network analysis is that not all relationships are of equal value. This is obvious from an examination of the many sociograms and social networks published in the literature: any given individual only grooms regularly with a proportion of the other members of its group (among many others, see network graphs given by Kummer 1968; Dunbar & Dunbar 1975; Voekl & Kasper 2009; Crofoot et al. 2011; Pasquaretta et al. 2014; Duboscq et al. 2016). Conventionally, the human literature distinguishes between *weak* and *strong* ties (Granovetter 1973; Sutcliffe et al. 2012) for this reason. Casual relationships often have a high turnover through time (for primate examples, see Altmann 1980; Dunbar & Dunbar 1988; Duboscq et al. 2016), as is the case in human social networks (Saramäki et al. 2014; Roberts & Dunbar 2015). The inclusion of all social contacts typically inflates the degree and, in the limit, leads unhelpfully to a saturated network (everyone grooms with everyone else) (James et al. 2009). A common strategy is, therefore, to divide ties between stronger more meaningful ones and weaker more casual ones by using some criterion based on the frequency of interaction. For present purposes, I follow previous studies (Dunbar 1984; Kudo & Dunbar 2001; Lehmann & Dunbar 2009a) and define a meaningful tie as one that accounts for at least 10% of an individual’s total social (i.e. grooming) effort (for justification and details, see *ESM*). This sets an upper limit on grooming clique size at 10 partners, but no species, or group, comes close to this. A tie that accounts for more than 10% of an individual’s social effort identifies a relationship likely to elicit coalitionary support (Dunbar 1984). I use undirected matrices (i.e. the data do not distinguish the direction of grooming), mainly because close relationships should be those that are reciprocated (i.e. each partner should invest equally in the relationship).

For each group, the grooming clique size of every adult in the group was first determined and then averaged, and then these were in turn averaged across all groups of the same species. Similarly, the size of all independent (i.e. non-overlapping) continuous network chains (n-cliques) in a group were first determined, averaged for each group, and then averaged across groups of the same species. Grooming clique size is independent of the actual size of group that animals happen to be in (Fig. S2), mainly because clique size is constrained by the species’ cognitive ability to handle relationships, and is thus characteristic of the species (Kudo & Dunbar 2001). Data on mean social group size for individual species are taken from (Campbell et al. 2008), the most recent comprehensive summary available, supplemented where necessary by more recent sources (see Table S1). Neocortex ratio, the best predictor of social group size (Dunbar & Shultz 2007) as well as of executive function cognition (Shultz & Dunbar 2010), is taken from Kudo & Dunbar (2001). Executive function cognition score is the arcsin transform of the mean proportion of correct trials on eight standard executive function tasks, the data for which were compiled from the published literature by Shultz & Dunbar (2010). Data on percent of day devoted to social grooming are taken from Lehmann et al. (2007).

In primates, species of the same genus typically exhibit very similar cognitive abilities and behaviour. To avoid inflating significance levels through phylogenetic autocorrelation, both structural indices and group size were averaged across species of the same genus to give mean values for individual genera and all analyses were carried out at genus level. Using generic means usually minimises the impact of phylogenetic effects. However, a genus-level analysis is essential for a second, more important, reason: phylogenetic methods remove grade differences, especially when these grades do not correspond to taxonomic divisions. Grades that cut across phylogeny are the focus of this analysis, and for this reason a genus-level analysis of raw data is necessary. This reduces the original sample from 34 species to 21 genera, although all results at genus level are also checked at species level (see *ESM*).

## Results

### Between-species comparison

Fig. 1 plots mean network size as a function of mean grooming clique size (number of principal grooming partners) for individual genera, and mean group size as a function of mean network size. There is a significant correlation in the case of network/clique (Fig. 1a: r=0.841, p<0.001, N=21), with a scaling ratio that does not differ significantly from 3 (t_19_[b=3]=0.03, p=0.979 (cf. Hill et al. 2008), but not in the case of group/network (Fig. 1b: r=0.303, p=0.181). However, a phylogenetic contrasts analysis (which removes the effect of grades) yields a highly significant relationship in the latter case (Fig. S3; contrasts in degree/clique, r=0.827, p=0.00008; contrasts in group/network, r=0.675, p=0.004), providing strong evidence that there is in fact an underlying relationship in this case too. The loss of significance when not controlling for phylogeny strongly implies that there is a grade effect. present Finally, values for individual species exhibit the same patterns, with similarly significant relationships in both cases (Fig. S4).

**Fig. 1.**
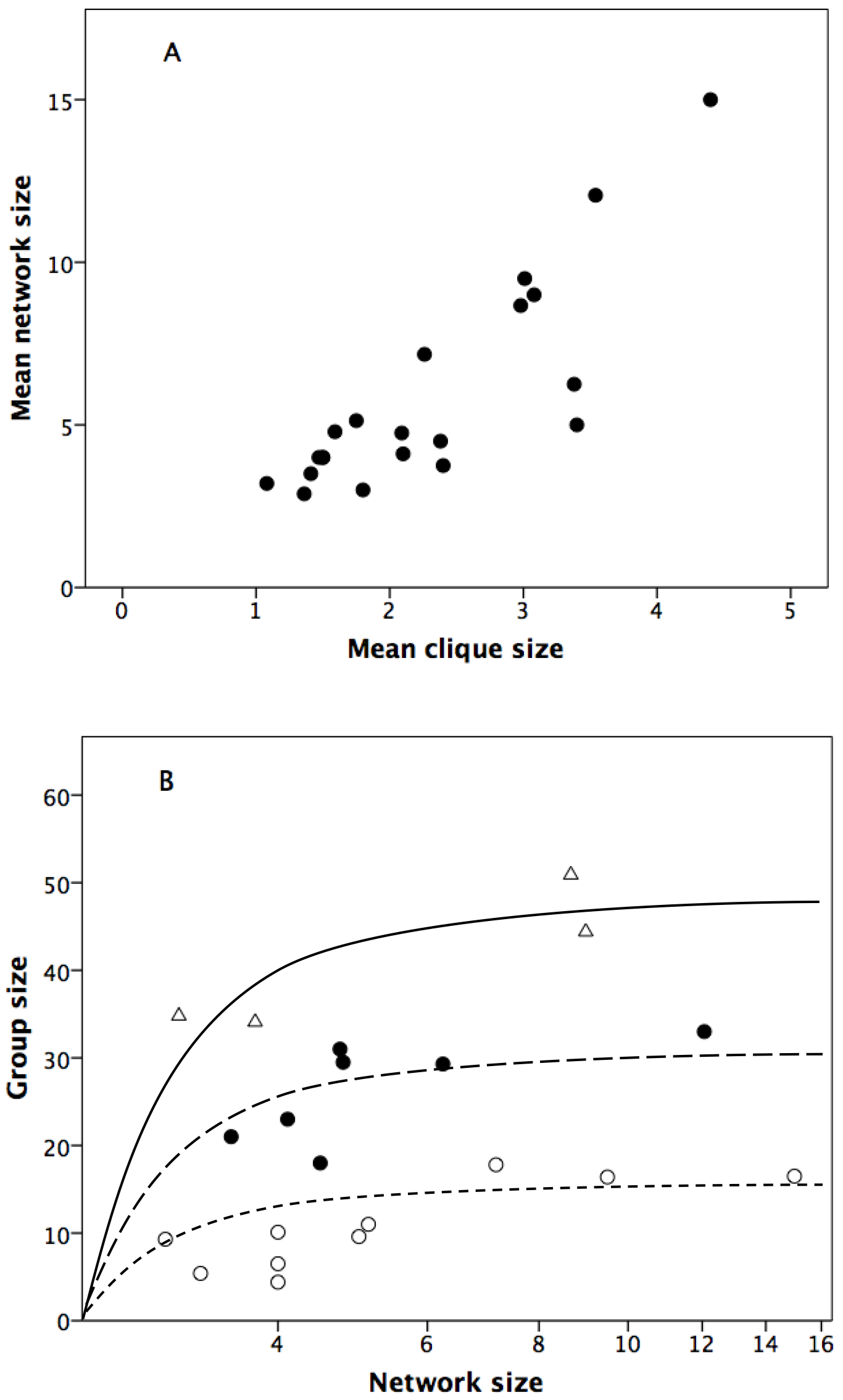
(a) Mean grooming network size plotted against mean grooming clique size and (b) mean group size plotted against log_10_ mean grooming network size for individual primate genera. Symbols in (b) indicate genera assigned to the three distinct social grades. Although the statistical analysis of the slopes for individual grades are based on simple linear regression, the regression lines in (b) are plotted on the assumption that they are in fact power curves anchored on the origin.

To explore this further, I examined residuals from the common RMA regression line in Fig. 1b. The residuals have an obviously trimodal distribution, unless one wants to consider a single outlier a grade in its own right (Fig. 2). A *k-means* cluster analysis of these residuals yields an optimal partition into three separate clusters (F_2,18_=93.3, p<0.0001), with both lower and higher order clustering yielding much poorer goodness-of-fits. Taken together, the three grades have individual slopes that are significantly positive (Fisher’s meta-analysis: *χ*^2^=22.65, df=6, p<0.001), with intercepts for the two upper grades differing significantly from that for the lower grade (t_12_=6.65, p<0.0001; t_15_=4.92, p=0.0002) but not from each other (t_9_=1.61, p=0.129), perhaps suggesting that the distinction between the upper and middle grades is less robust than that between them and the lower grade. A multiple regression of group size against network size with grade as a dummy variable yields a significant regression with a radically improved goodness of fit (with grades, r^2^=0.919 vs r^2^=0.092 without grades; with grades, F_2,18_=102.1, p<0.00001), and significant partial regressions for both network size (t_p_=4.51, p=0.0003) and grade (t_p_=13.56, p<0.0001).

**Fig. 2.**
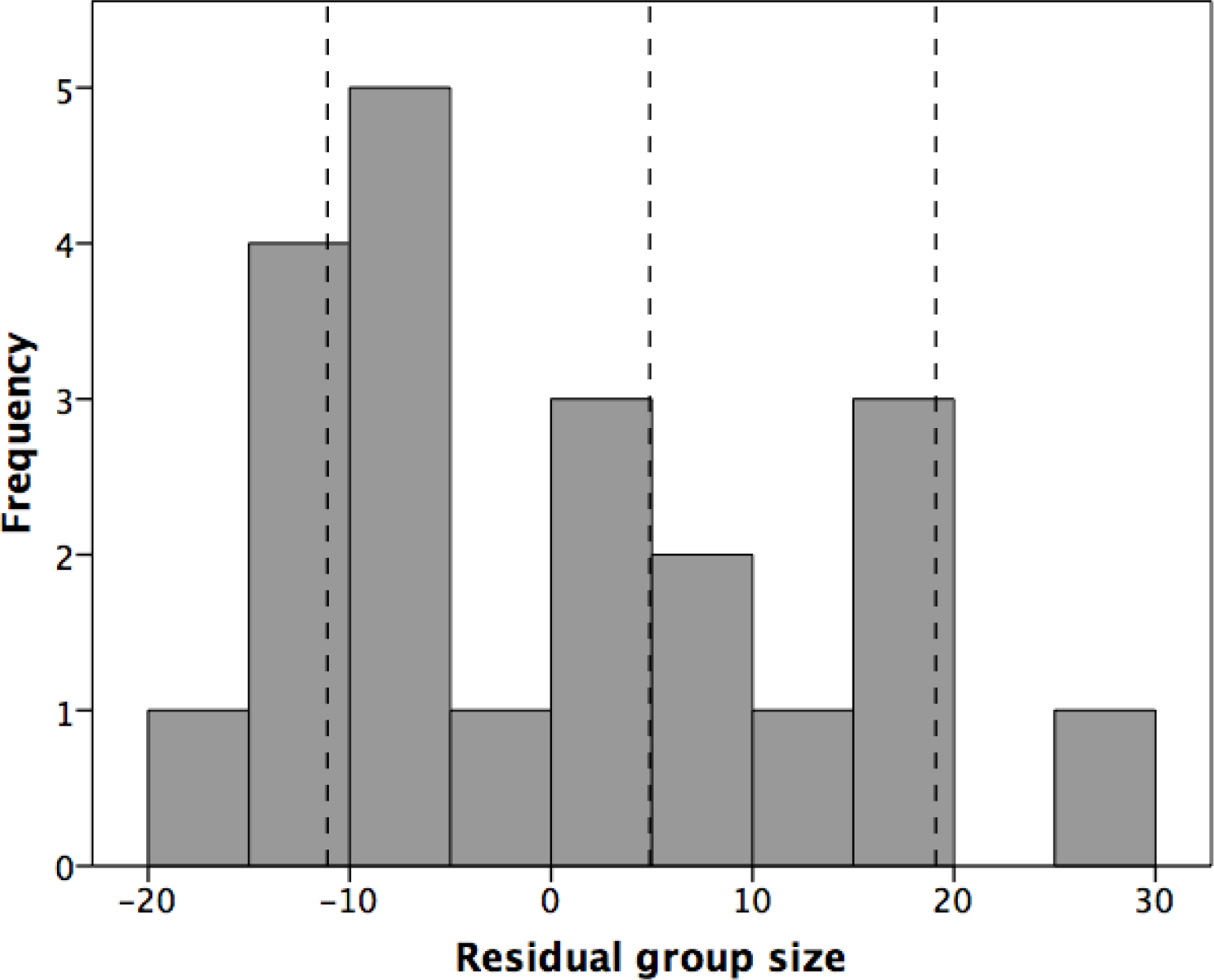
Distribution of residuals from the common regression line for the genera in Fig. 1(b).

The three grades in Fig. 1b include, left to right:

Upper grade: *Erythrocebus* [1], *Procolobus* [1], *Papio* [1,1], *Pan* [1,1]
Middle grade: *Chlorocebus* [1], *Saimiri* [1], *Sajapus* [1], *Ateles* [1], *Theropithecus* [1]*, Semnopithecus* [1], *Macaca* [1,1,1,0]
Lower grade: *Colobus* [0], *Propithecus* [0], *Alouatta* [1], *Indri* [0], *Callicebus* [0], *Eulemur* [1], *Trachypithecus* [0,0], *Cercopithecus* [0,0,0]*, Cebus* [1], *Presbytis* [0], *Lemur* [1]

The network:clique ratios for individual species approximate 3, and do not differ across the three grades (Fig. 3a: F_2,30_=0.69, p=0.510). However, the group:network ratios differ significantly across grades (Fig. 3b: F_2,29_=15.3, p<0.001), with the three grades differing significantly from each other (Scheffé post hoc tests: p≤0.044). Mean network:clique ratios are 2.71, 2.95 and 2.53, while mean group:network ratios are 2.31, 4.23 and 7.02, respectively, for the three grades.

**Fig. 3.**
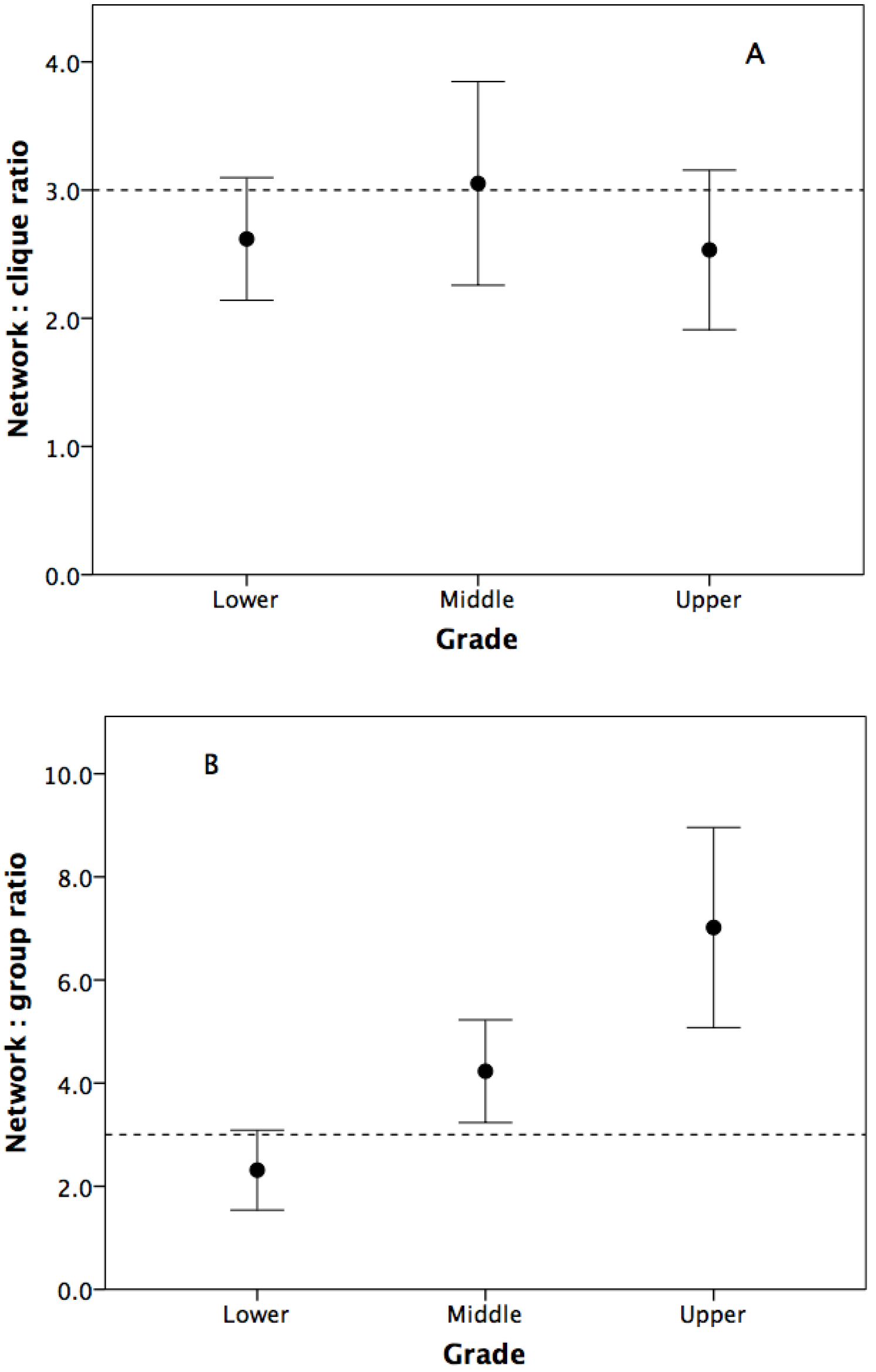
Mean (±2 se) (a) network:clique ratio and (b) group: network ratio for the three grades shown in Fig. 1b. The dashed line marks a scaling ratio of 3.

### Cognition, time budgets and social structure

The three grades differ significantly in neocortex ratio (Fig. 4: means of 1.78±0.55 vs 2.37±0.16 vs 2.78±0.43; F_2,18_=8.79, p=0.002). Post hoc tests confirm that the lower grade differs significantly from the other two grades (Scheffé tests: p=0.041 and p=0.004), but the two upper grades do not differ significantly from each other (p=0.343). The three grades also differ significantly in executive function cognitive competences (Fig. 5: F_2,15_=8.32, p=0.004), with the lower grade being significantly different from the two upper grades (Scheffé tests: p=0.049 and p=0.004), but the two upper grades again not differing significantly (p=0.105).

**Fig. 4.**
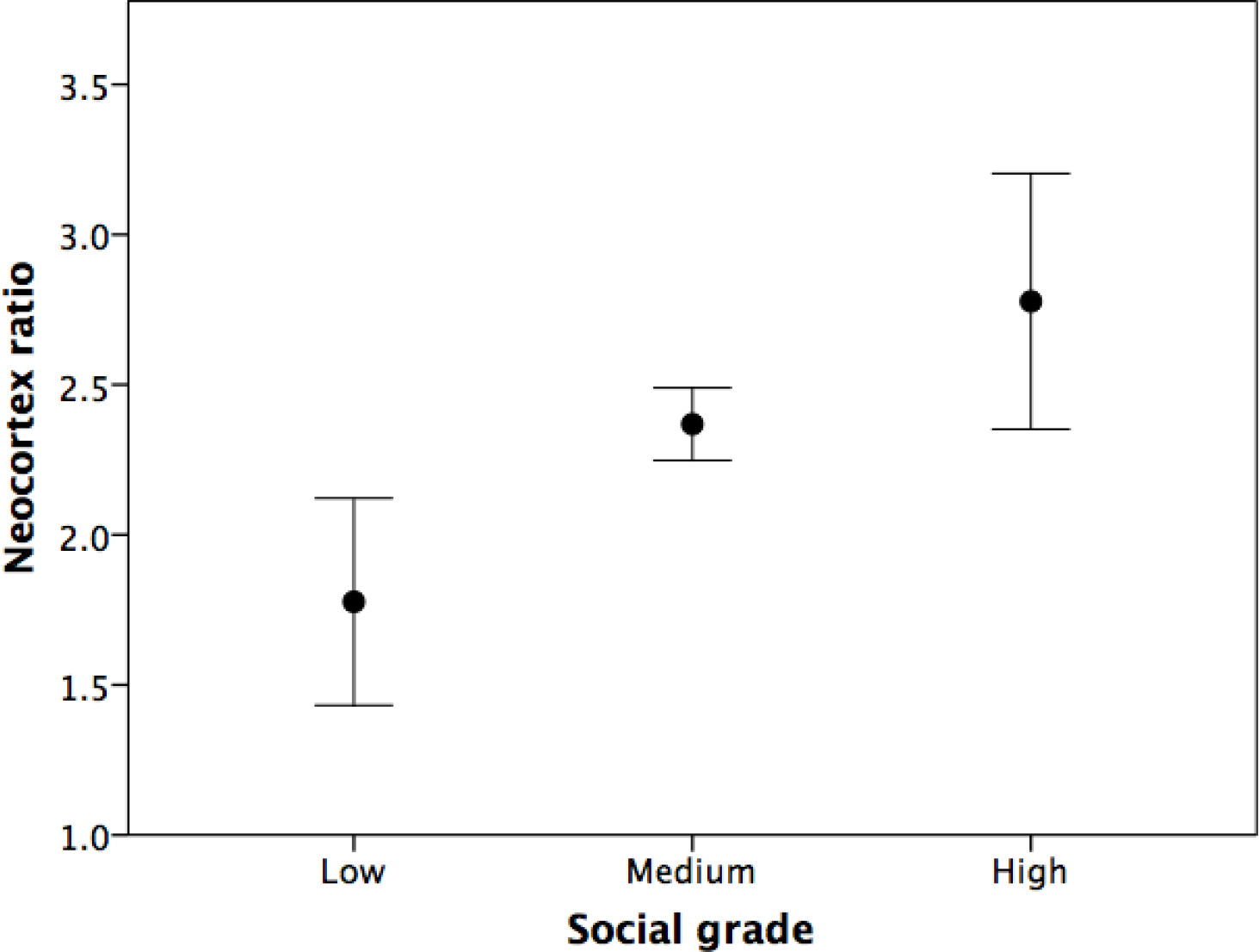
Mean (+2 se) neocortex ratio for the three social grade shown in Fig. 1b.

**Fig. 5.**
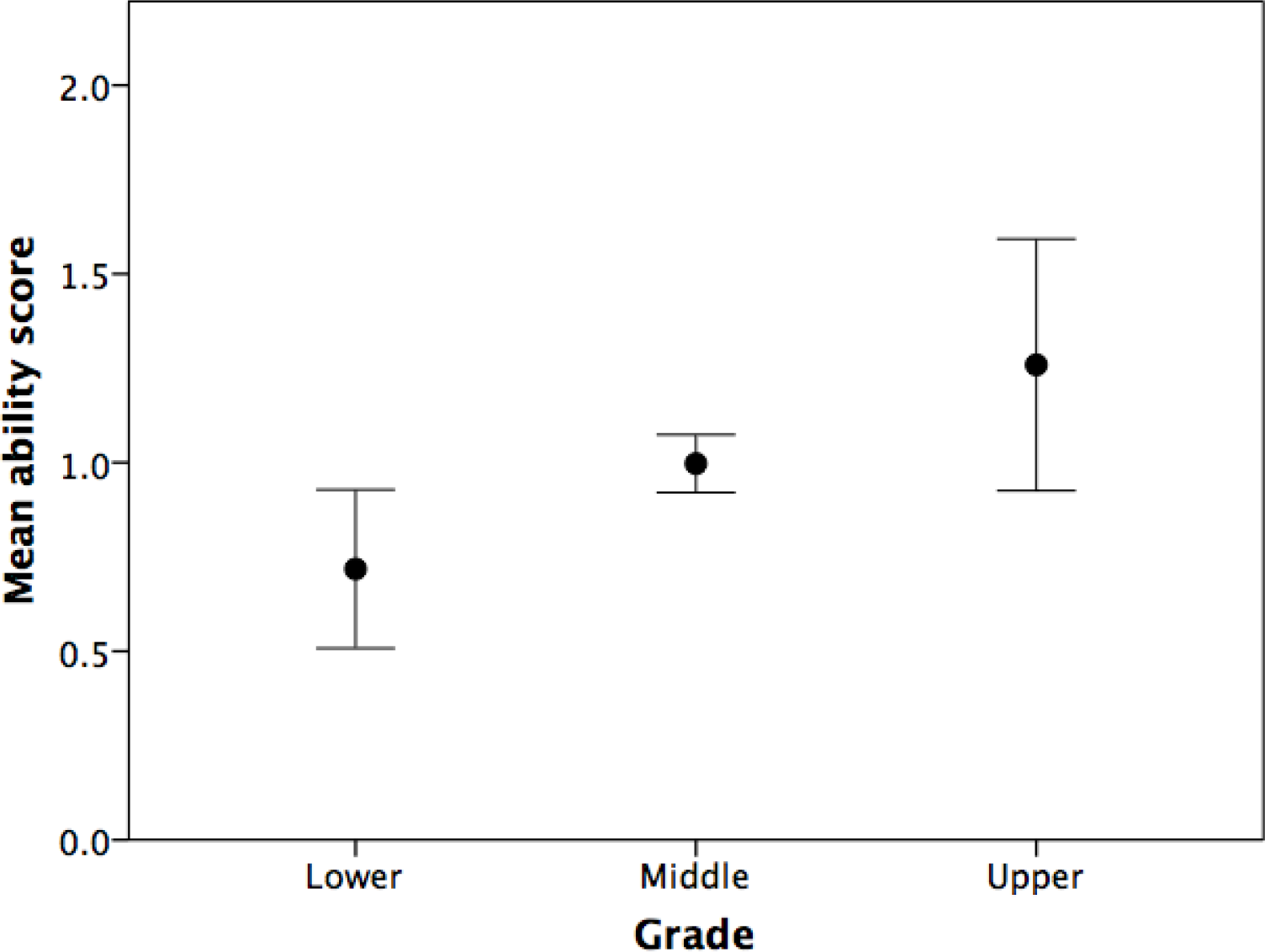
Mean (±2 se) executive function ability score for individual species in the three grades. Ability score is the arcsin transform of the proportion of correct trials on 8 standard executive function tasks. Data from (8).

While there is, overall, a significant positive correlation between species mean group size and mean percent of the day devoted to social grooming (Fig. 6: F_1,20_=7.47, p=0.013) (cf. Dunbar 1991; Lehmann et al. 2007; Dunbar & Lehmann 2013), the data in fact partition into two distinct grades with parallel slopes. However, this obscures a more complex relationship. A backwards stepwise multiple regression with mean species grooming time as the dependent variable and mean group size, mean network size, mean clique size and grade (as a dichotomous dummy variable: lower versus middle+upper) as independent variables yields a highly significant model (F_3,18_=5.59, p=0.006) with a negative relationship with clique size (p=0.035), a positive relationship with network size (p=0.005), and a significant effect of grade (p=0.007), and no effect due to group size (Table 1).

**Fig. 6.**
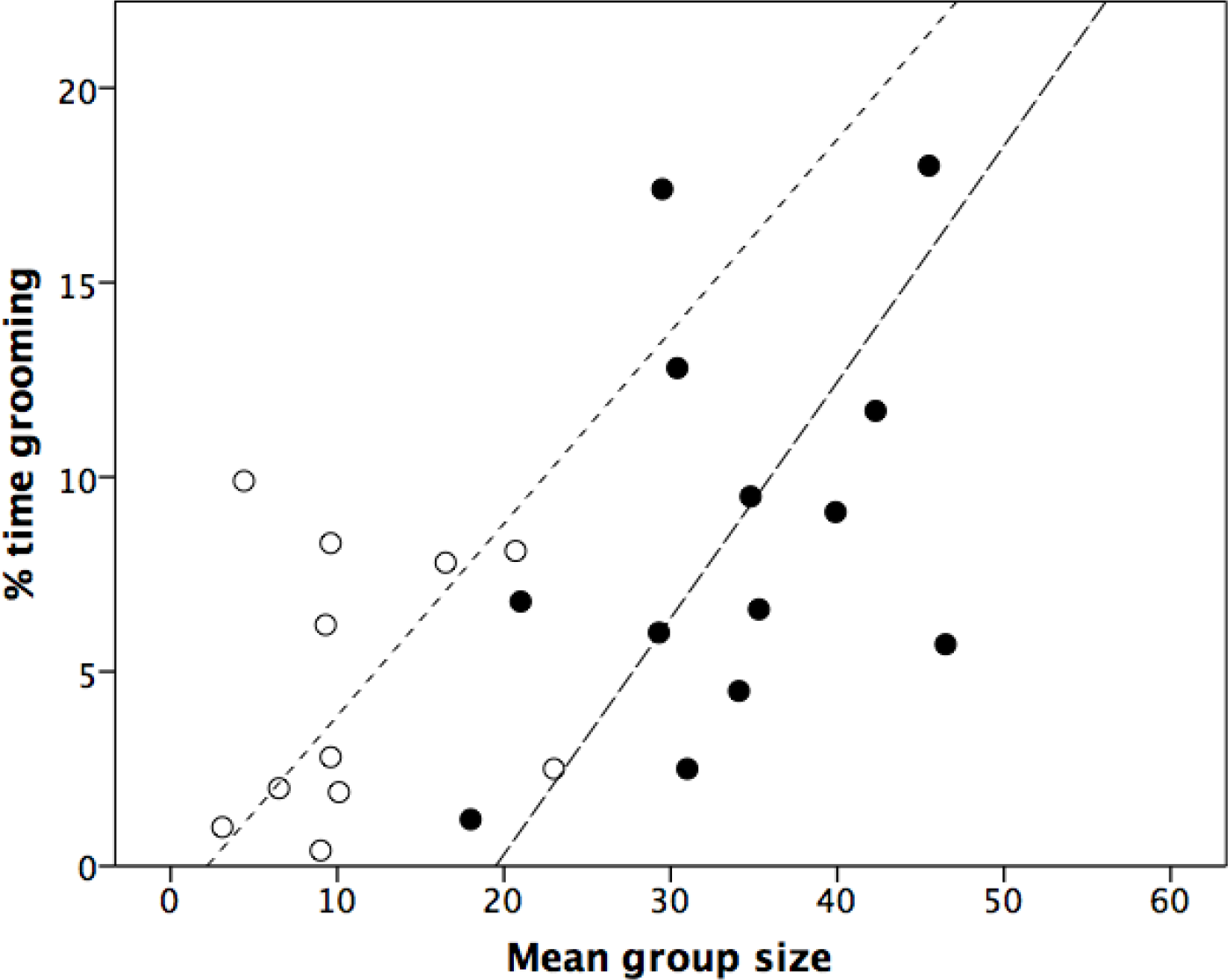
Mean percentage of daytime devoted to social grooming by individual species, plotted against species mean social group size. Species are distinguished according to whether their genera are classed as upper grade (solid symbols) or lower grade (open symbols) in Fig. 1b. Regression lines are LSR regressions (sold line: upper grade; dashed line: lower grade).

**Table 1.** Best fit backward stepwise multiple regression analysis of percent time spent grooming as the dependent variable.

| 540 | Variable | Slope | Standardized slope | t | p |
| --- | --- | --- | --- | --- | --- |
| 541 | ----- |  |  |  |  |
| 542 | Constant | 0.469 |  |  |  |
| 543 | Clique size | -4.218 | -0.830 | -2.28 | 0.035 |
| 544 | Network size | 1.135 | 1.108 | 3.16 | 0.005 |
| 545 | Grade* | 5.609 | 1.835 | 3.06 | 0.007 |

Excluded variable: group size
\* lower grade taxa scored 1; middle and upper grade taxa scored 2

## Discussion

The results suggest that group-living primates divide into at least two, possibly three, grades of sociality in terms of the relationship between grooming network size and group size. Species from the lower social grade are heavily restricted in the range of group sizes they can have, but the more social species are somehow able to maintain larger groups for a given grooming clique size. For many lower grade taxa, group size and network size are synonymous, but the two higher grades are able to maintain groups that are 4-6 times the size of their grooming networks. In other words, the groups of these species seem to contain a large number of effectively unconnected (or weakly connected) grooming networks that are likely to constitute potential fracture points for group fission. Nonetheless, even though the quality of a dyadic relationship is related to the amount of effort directly invested in it for both monkeys (Dunbar 1980; Seyfarth & Cheney 1984) and humans (Sutcliffe et al. 2012), upper grade species are seemingly able to maintain group cohesion without having to groom with everyone in the group, whereas lower grade species seemingly cannot.

Fig. 1a suggests that there is an upper limit at around five individuals that can be maintained as grooming partners. This is identical to the number of intimate social partners found in humans (mean≈5: Sutcliffe et al. 2012; Burton-Chellew & Dunbar 2015). In primates, the grooming clique functions as the principal defence coalition: individuals in this category are more likely to come to each other’s aid when one of them is challenged (Dunbar 1980, 1984; Seyfarth & Cheney 1984). This set of grooming partners also provides the basis for an extended grooming network (or *n*-clique, the number of individuals that can be reached directly or indirectly through a continuous chain) that is three times larger, and this is true of all the genera in the sample. Hill et al. (2008) found a scaling ratio of ~3 for grouping levels in a sample of mammals (including three primate species) that live in multilevel societies, and a similar layering with a scaling ratio of ~3 has been widely documented in human social networks (Hill & Dunbar 2003; Zhou et al. 2005; Hamilton et al. 2007; Dunbar et al. 2015; MacCarron et al. 2016). The larger sample provided by this study however, suggests that there may be a distinction between lower grade taxa whose networks more or less correspond to their groups and higher grade taxa (those sampled by Hill et al. 2009) which have much higher group:network scaling ratios such that their groups consist of several weakly connected sub-networks.

These results suggest that upper grade taxa can somehow maintain temporal coherence between sub-groups that do not interact directly very often, whereas lower grade taxa cannot. It is noteworthy that the upper grade taxa differ from the middle grade mainly in the fact that all four of the upper grade genera either occupy terrestrial (and therefore predator-risky) habitats (Lehmann & Dunbar 2009b; Bettridge & Dunbar 2012) or typically experience unusually high predation risk, notably from chimpanzees (e.g. *Procolobus*: Stanford 1995). It seems that these species are either being pushed beyond their capacities in order to live where they do or have evolved additional cognitive capacities to cope with larger groups (as may be the case in baboons and chimpanzees).

The capacity to maintain social cohesion without grooming with everyone appears to depend on increased neocortex volume, and not phylogeny, and may thus depend on novel aspects of cognition. Indeed, in terms of executive function, there are correlated differences between the grades in cognitive ability. The grooming data (Fig. 6) suggest that while lower grade taxa simply increase grooming time to allow a larger clique (and hence group) size, taxa in the upper grades seem to trade off investment in their close allies (principal grooming partners) to allow time for more casual interactions with other members of their extended network in a way that probably gives these extended networks greater coherence and structural stability. However, upper grade taxa are also managing to do something else, and that is to ensure that several other similar network subgroups remain within their group despite the stresses imposed by having more individuals in the group and despite not grooming with them. In part, this might be because individuals are able to acquire knowledge about third parties with whom they do not interact directly by observing them interacting with other members of the group, and later using this knowledge to infer, for example, where an individual sits in the wider dominance hierarchy (Bergman et al. 2003; Schino et al. 2006), whom they have alliances with (Datta 1999; Cheney & Seyfarth 1999) and, on the basis of observed reputation, how trustworthy they might be (Silk 1999; Russell et al. 2008). Although competences of this kind have often been documented in upper grade taxa, evidence for these capacities is, so far at least, largely absent from any lower grade genus.

As Fig. 4 suggests, the structural differences in the way these species’ social groups are organised relate to the relative sizes of their brains, in particular the size of the neocortex, and Fig. 5 shows that there are correlated differences in cognitive ability, suggesting these may play an important role in facilitating the higher order competences that underpin the upper grade form(s) of sociality. (Note that these executive function competences have been shown to correlate with neocortex volume across primates, even when correcting for phylogenetic relationships: Shultz & Dunbar 2010.) The frontal lobes play a crucial role in managing social relationships, and hence in individual differences in personal ego-centric network size in both humans (Lewis et al. 2011; Powell et al. 2012; Kanai et al. 2012) and macaques (Sallet et al. 2013), as well as social cognition (such as the capacity to integrate others’ perspectives with one’s own: Powell et al. 2010, 2014). Brodman area 10 in the frontal pole of the neocortex, which is found *only* in anthropoid primates, seems to be crucial for allowing animals to engage in the kinds of advanced forms of executive function cognition (notably one-trial learning, causal reasoning, strategic comparisons and inhibition: Passingham & Wise 2012) that are likely to be essential for the management of complex social relationships. It may be relevant that no non-primate mammals that lives in primate-like bonded (Silk et al. 2010a; Dunbar & Shultz 2010; Massen et al. 2010) social groups (e.g. equids, canids, tylopods, hyaenids) (Shultz & Dunbar 2010) has a mean group size that exceeds 30 animals; the only exceptions are elephants and delphinids, both of whom have neocortices in the cercopithecine primate range as well as primate-like multilevel social systems (Hill et al. 2008). Clearly, more data on the cognitive and social behaviour of other mammal and primate species are needed in order to explore this further.

The grooming data of Fig. 6 suggest (i) that animals adjust their grooming time commitments mainly to the size of their social networks, rather than group size as a whole, (ii) that proportionately more time is committed to grooming in those species where groups are multilevel-structured, and (iii) that they free off time to make this possible by reducing their investment in their core grooming partners (clique members). It seems that the limit on the number of grooming partners that an individual can have (its degree) is ultimately set by the amount of time it can afford to devote to social interaction, and this in turn is likely to be determined by the extent to which its diet imposes demands on foraging and resting time (Dunbar et al. 2009). The lower grade taxa differ from those in the upper grades by being more folivorous (35.5±26.9% leaf in diet vs 18.1±15.4% for upper grade taxa; data in Korstjens et al. 2010). Species that depend on hindgut fermentation to extract nutrients from high-fibre leaf-based diets are obliged to devote significant amounts of time to resting in order to allow fermentation to occur because the heat generated by any form of activity suppresses the gut flora responsible for fermentation (van Soest 1982). This phenomenon is familiar in ruminants, who are obliged to lie down and rest when ruminating (van Soest 1982). This severely limits the time available for social interaction in the more folivorous primates (Dunbar 1988; Korstjens & Dunbar 2008), and seems to be mainly responsible for their smaller, less structured groups.

A switch from a more leaf-based diet to a more fruit-based one has been shown to be responsible for the contrast in social group size between *Colobus* (a member of the lower grade, mean group size = 9.5; leaf = 59% of diet) and *Procolobus* (a member of the more intensely social upper grade, mean group size = 33.8; leaf = 46%): *Colobus* populations spend an average of 62.4% of their day resting, compared to 44.8% for *Procolobus* populations (Korstjens & Dunbar 2008), and thus have considerably less time available for social grooming, which in turn limits group size. Giving *Colobus* species a *Procolobus*-like diet allows them to live in groups as large as those found in *Procolobus*, and *vice versa* (Korstjens & Dunbar 2008). The issue, then, is that improvements in diet quality allow less time to be spent foraging, thereby freeing off time for social interaction (and hence bonding larger groups) as well as allowing the higher nutrient throughput needed for larger brains to be evolved to handle more complex relationships (Dunbar & Shultz 2017).

In sum, the apparent ability to support a larger group with less investment seems most likely to involve a phase shift in socio-cognitive competences, probably dependent on significant increases in brain size that allow novel behaviours (Shultz & Dunbar 2010). One way this might have been achieved is by enabling individuals to maintain several qualitatively different kinds of relationships at the same time in a way reminiscent of Granovetter’s (1973) distinction between weak and strong ties in human networks. Whether primates as a whole represent just two or more than two such phase shifts remains to be investigated in more detail.

## Acknowledgments

My research is supported by a European Research Council Advanced Investigator grant.

## Ethics statement

This study did not require ethical approval.

## Data accessibility

The data can be found in the ESM.

## Authors’ contributions

The named author is the sole author.

